# A macromutation eliminates colour patterning in captive butterflies

**DOI:** 10.1101/2021.10.29.466422

**Authors:** Joseph J. Hanly, Luca Livraghi, Christa Heryanto, W. Owen McMillan, Chris D. Jiggins, Lawrence E. Gilbert, Arnaud Martin

## Abstract

Captive populations often harbor variation that is not present in the wild due to artificial selection. Recent efforts to map this variation have provided insights into the genetic and molecular basis of variation. *Heliconius* butterflies display a large array of pattern variants in the wild and the genetic basis of these patterns has been well-described. Here we sought to identify the genetic basis of an unusual pattern variant that is instead found in captivity, the *ivory* mutant, in which all scales on both the wings and body become white or yellow. Using a combination of autozygosity mapping and coverage analysis from 37 captive individuals, we identify a 78kb deletion at the *cortex* wing patterning locus as the *ivory* mutation. This deletion is undetected among 458 wild *Heliconius* genomes samples, and its dosage explains both homozygous and heterozygous *ivory* phenotypes found in captivity. The deletion spans a large 5’ region of the *cortex* gene that includes a facultative 5’UTR exon detected in larval wing disk transcriptomes. CRISPR mutagenesis of this exon replicates the wing phenotypes from coding knock-outs of *cortex*, consistent with a functional role of *ivory-*deleted elements in establishing scale color fate. Population demographics reveal that the stock giving rise to the ivory mutant has a mixed origin from across the wild range of *H. melpomene*, and supports a scenario where the *ivory* mutation occurred after the introduction of *cortex* haplotypes from Ecuador. Homozygotes for the *ivory* deletion are inviable, joining 40 other examples of allelic variants that provide heterozygous advantage in animal populations under artificial selection by fanciers and breeders. Finally, our results highlight the promise of autozygosity and association mapping for identifying the genetic basis of aberrant mutations in captive insect populations.

## Introduction

Captive populations can harbour variation that is not observed in the wild. Aside from domestication for agricultural or commercial yield, animals and plants have often been selectively bred for ‘beautiful’, ‘interesting’ or ‘unnatural’ aesthetic traits, especially colour and pattern variants (1,2). In recent decades, the genetic basis of artificially selected variation has begun to be mapped in many organisms used in agriculture, floriculture and the pet trade, including the genetic mapping of a large number of genes responsible for flower colour variation (3,4), melanin-based coat colour in domestic mammals (1), plumage patterns and colours in birds (5,6), and chromatophore distribution in squamates and fishes (7). These variants can affect genes involved in the enzymatic production, transport and deposition of pigments like the melanin and carotenoid pathways (8,9), or can be caused by genes that affect signalling or cell type differentiation (as with chromatophore distribution in squamates(10)). The extensive set of artificial and domesticated genetic variants has been used repeatedly both for studies of genotype-to-phenotype relationship, and for understanding the genetics of disease states in humans (11).

Notably, several cases have been identified where a gene underlying an artificially selected variant is also responsible for natural variation in other populations or species. This is the case with *BCO2*, where an artificially selected protein coding variant in cattle affects milk colour (12), and a wild type regulatory variant in wall lizards causes changes to ventral scale colour (13). The gene Agouti has repeatedly been mapped in cases of pigment differences, both in cases of domestication [*e*.*g*. in the chicken (14), horse (15), and rabbit (16)], and in natural variation (*e*.*g*. in warblers (17), humans (18) and snowshoe hare (19)). This reuse of hotspot genes in wild and captive populations can even be observed at the within-species level; multiple separate variants in agouti have been linked to pigment variation in sheep - both in wild and captive populations, and caused by protein coding mutations, *cis-*regulatory mutations and copy number variation (20,21). As such, studies of domesticated variation can inform our understanding of natural variation too, both on macro- and micro-evolutionary scales

As the primary selective force in captive populations is the eye of the selector, one may often observe variation that is not seen in nature because it is not optimally fit (22–25). Fanciers and breeders have purposefully selected for variation that causes deleterious effects in combination with colour and pattern differences, applying a regime of selection that favours the phenotype of interest over fitness. This applies especially to distinctive colouration, which can be straightforward to observe and maintain in a captive stock. Examples include coat color variants for Merle dogs, where homozygotes for a retrotransposon insertion in the *SILV* gene have an increased risk of deafness and blindness (26,27), and in overo horses, where foals homozygous for a mutated Endothelin Receptor B (*EDNRB)* develop Lethal White Syndrome (28). “Lemon frost” geckos have been selected for their unique colour, but exhibit an increased risk of iridophoroma, the formation of tumours from iridophores (analogous to melanoma) (7). Such reduced fitness has also been identified in floriculture, where white petunias with mutations to the *AN1* gene have deficient vacuole acidification and a weakened seed coat (29), though an artificially selected mutation to the same gene causing a similar effect in the morning glory does not appear to carry the same deleterious fitness effects (3). Whether they are the direct targets of selective breeding, or collateral effects of inbreeding and bottlenecks, deleterious mutations that would be purged by natural selection in the wild are common in captive populations (30).

The majority of work mapping artificially selected or domesticated variation in animals has taken place in vertebrates, with the notable exceptions of the economically important silkworm (31). *Heliconius* butterflies, a model system for the genetic study of color pattern adaptations in the wild (32,33), have also been maintained in captivity by butterfly breeders continuously since at least the mid-20th century, including at notable tourist attractions like Butterfly World in Florida. There, they have been selectively bred for hundreds of generations with special attention paid to pattern variations that deviate from phenotypes normally observed in the wild. Over an extended period of inbreeding, selection and occasional out-crossing, novel color variations have appeared in this captive population, including the ‘Piano Keys’ (PK) pattern, and a deleterious mutation that occurred within the PK genetic stock, here dubbed *ivory* (Fig 1).

**Fig 1.**
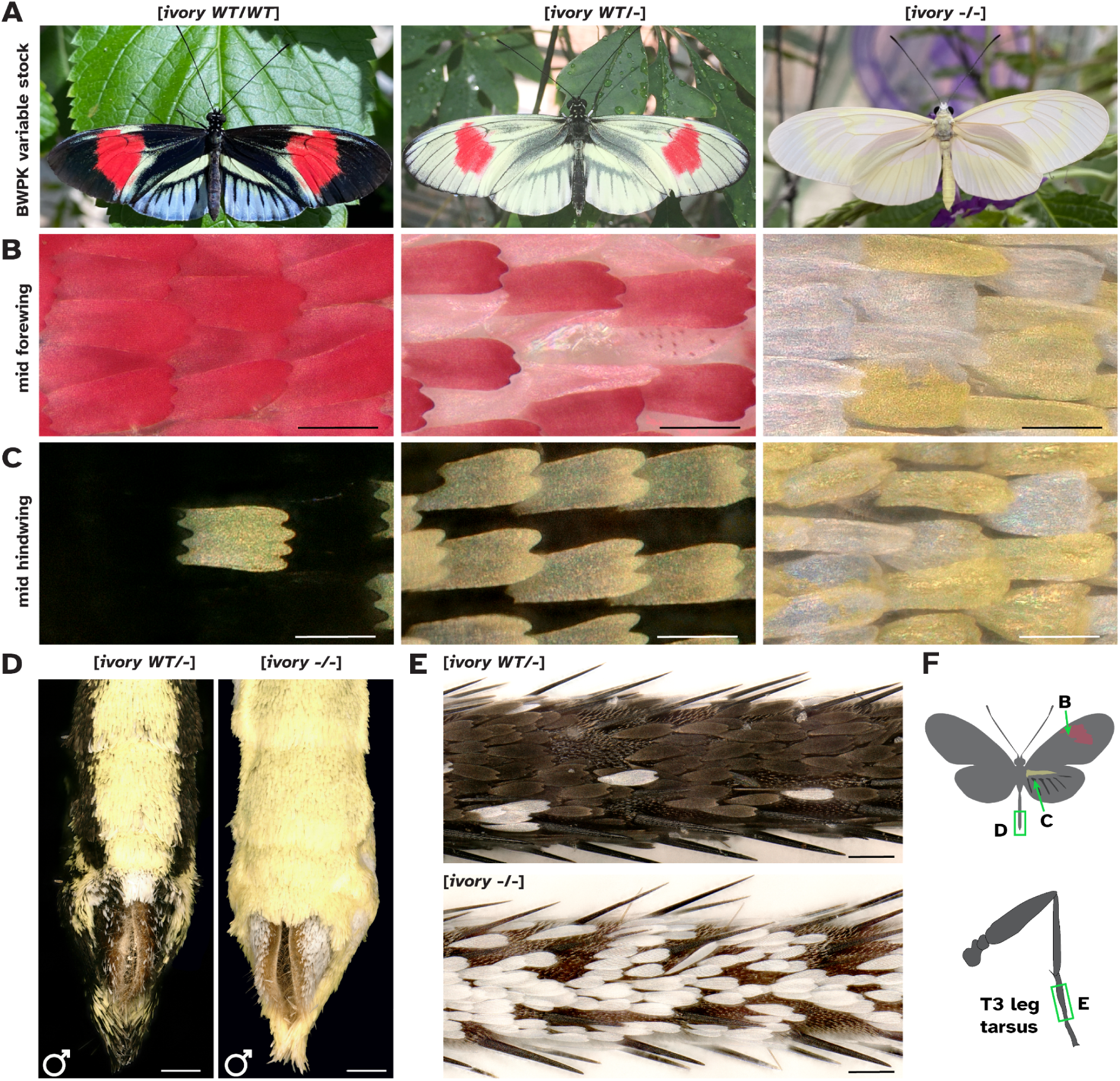
Phenotypes of *H. melpomene* BWPK butterflies. **(A)** Ivory phenotypes in the *H. melpomene* “Piano Keys” in the UT Austin stock (BWPK). The three color states (from left to right “Dark BWPK”, “Pale BWPK”, “ivory”) depend on the allelic dosage of a co-dominant mutation. Ivory homozygotes are inviable and only found among offspring from two pale heterozygotes. **(B)** Magnified view of the forewing red band region. In the *[ivory WT/-*] state, abnormal scales have formed in the red region, with red pigment in granules and scales curled, while in [*ivory -/-*], all scales are yellow or white. **(C)** Magnified view of a central hindwing region that is black in [*ivory WT/WT*]. The [*ivory WT/-*] wing has all cover scales as yellow or white while all ground scales remain black. All scales are yellow or white in [*ivory -/-*]. **(D-E)** Complete replacement of melanic scales by yellow-white scales in the abdomen **(D)** and legs **(E)** from [*ivory -/-*] homozygotes. **(F)** Cartoon of image locations for B-E. Scale bars : **B-C** = 50 μm ; **D** = 500 μm ; **E** = 100 μm.

The genetics of wing patterning have been extensively studied in *Heliconius* (34). Linkage and association mapping have pinpointed regulatory regions around a small toolkit of genes as being responsible for much of the pattern variation seen in wild populations, including the transcription factor *optix* (35–38), the signaling ligand *WntA* (39,40), and *cortex*, a *cdc20* homolog with a currently-unidentified function (41,42). We used whole genome resequencing, association mapping and autozygosity mapping to determine that the *ivory* mutation is caused by a large deletion at the patterning gene *cortex*, which likely occurred *de novo* in the captive population.

## Methods

### Stock history

The captive population of *Heliconius melpomene* was initially by JRG Turner and others from around Central and South America, and were transferred from his genetic research stocks at the University of Leeds to the former London Butterfly House (LBH) initiated by Clive Farrell 1981 as a multi-race hybrid population. They were then acquired from Tom Fox of LBH by R. Boender at the MetaScience Butterfly Farm in Florida about 1985. These hybrid stocks formed a core of Boender’s inhouse hybrid *Heliconius* display stock when Butterfly World (BW) opened in 1988. Only one known introduction of wild-caught *H. melpomene* occurred to this stock, of *H. m. cythera* collected in the early 1980’s by R. Boender and T. Emmel around Tinalandia, Ecuador. After this introduction, Boender began selection for a novel pattern that is termed “Piano Key” in this paper. The exact mix of races that were transferred from LBH to BW are not known, but contain pattern alleles that are characteristic of *H. melpomene* races of Suriname and the Guianas (LEG, personal obs).

Since the Piano Key phenotype was first noticed by R. Boender over 30 years ago, the Piano Key stock population has been maintained in separate compartments for around 400 generations at Butterfly World and has long been fixed for the PK phenotype (RB, personal communication). Thus, we referred to it as *H. melpomene* BWPK throughout. Within the last decade, novel mutant butterflies appeared with fewer melanic scales, initially termed “pale PK”. These were subsequently separated to form a selected sub-population at BW. Soon after, virtually pure white butterflies incapable of flight began appearing in this selected stock, herein referred to as *ivory*. In 2015 “Pale PK” were sent to UT Austin and maintained by LEG in climate-controlled greenhouses (see S1 Table for phenotypes and collection dates) to investigate the genetics of this mutation. We now know that “Pale PK” phenotypes are heterozygous for ivory mutation and the pure white and flight-disabled phenotypes are homozygous for the ivory mutation (see below).

### Imaging

Pinned specimens were imaged with a Nikon D5300 camera mounted with a Micro-NIKKOR 105mm f/2.8 lens. Scale phenotypes were imaged with a Keyence VHX-5000 microscope mounted with a VH-Z100R lens.

### Short read DNA sequencing

DNA was extracted from thoracic tissues of 37 *H. m*. BWPK individuals using the Qiagen DNeasy Blood & Tissue Kit, RNAse-treated, and used to prepare a multiplexed sequencing library with the TruSeq PCR-free DNA protocol. Samples were sequenced on an Illumina NovaSeq S1 150bp PE run, yielding 13x mean coverage per individual. Sequencing reads are accessible via NCBI SRA under the project accession number PRJNA663300.

### Genomic analyses

Samples were aligned to Hmel2.5 (43) retrieved from LepBase (44) with BWA-MEM using default parameters (45), and variants called using GATK v4.1 with tools HaplotypeCaller and GenotypeGVCFs using default parameters (46). Variant sites were accepted if they were biallelic and the quality (QUAL) value was ≥30. SNPs were phased with Beagle 4.1 (47). Phased SNP variants were used to perform principal component analysis (PCA) using the Eigensoft module SmartPCA(48). SNP association was carried out in PLINK v.1.9, with 1000 permutations(49). For phylogenetic analyses, we followed the TWISST pipeline(50); briefly, data were phased in Beagle 4.1 with a window size of 10,000 and overlap of 1000. Trees were built from windows of 50 SNPs, and then analysed with TWISST using the five groups “East”, “West”, “Ecuador”, “Atlantic”, “BWPK” (S2 Table).

#### *De novo* assembly for indel breakpoints

To find precise indel breakpoints, a subset of samples were *de novo* assembled with Velvet (51) using default parameters. The resulting contigs were searched with BLAST (52) for the genomic region including the deletion. Scaffolds that bridged the indel breakpoints were selected, and aligned to the Hmel2.5 reference genome with MAFFT (53).

### CRISPR/Cas9 mutagenesis

We designed sgRNAs against the putative *H. erato cortex* promoter corresponding to N_20_NGG sites within the distal promoter/5’UTR. In order to mutagenize the locus, we designed three sgRNAs (Table 1) using the “find CRISPR sites” algorithm within the Geneious software (https://www.geneious.com/). Guide specificity and off-target effects were assessed by scoring against the *H. erato* reference genome. sgRNAs displaying low off-target scores were then synthesised commercially by Synthego, and mixed with Cas9 at a concentration of 500:500 ng/ul respectively. Embryonic injections were performed as previously described (42) within 1-3 hours after egg laying, after which larvae were allowed to develop on a diet of *P. biflora* until adult emergence.

**Table 1.**
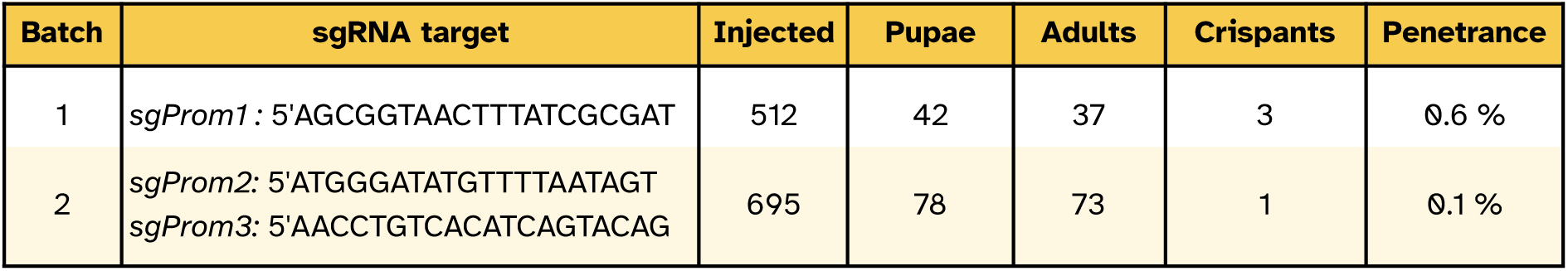
CRISPR mutagenesis of the *cortex* distal promoter/5’UTR in *H. erato*.

## Results

### *H. melpomene* BWPK - artificially selected wing patterns in an insectary stock

The stock maintained at Butterfly World in Florida and at UT Austin is here named *H. melpomene* BWPK (see Methods - Stock history section). The insectary-bred line is polymorphic for several wing pattern elements found in wild populations, including the red pattern elements (Fig 1). Additionally, *H. m*. BWPK includes aberrant pattern features not observed in natural populations. First, in the pattern dubbed ‘Piano Keys’ (Fig 1A), white or yellow hindwing marginal elements extend along the veins towards the discal cell. While distal yellow hindwing pattern elements are observed in some wild pattern forms of *Heliconius*, including *H. melpomene cythera* from Ecuador and *H. cydno*, the extent of these marginal elements in *H. m. BWPK* is much greater than any of these, occurring over most of the area of the hindwing except for the intervein regions that would otherwise be taken by the red hindwing rays.

Occasional butterflies with more extensive regions of yellow or white scales were identified (Fig. 1A centre). Other than the wing pattern differences, both forms of butterflies exhibit typical feeding, mating, egg-laying and flight behaviours to other insectary-reared *Heliconius* (*e*.*g*. S1 Movie). However, the offspring of a mating of two pale *H. m*. BWPK will include completely white-winged and -bodied butterflies (Fig 1C). We dub these butterflies ‘ivory’ in reference to their emergence in the ‘Piano Keys’ stock, and inferred that the pale morph of *H*.*m*. BWPK represents a heterozygous state for a mutation that causes the ivory phenotype in the homozygous state.

The ivory butterflies have melanin pigments in their body cuticle and eyes, indicating that their capacity to synthesize and deposit melanin has not been perturbed. However, all scales on the wings and body are white or yellow, with no black or red scales, with the exception of a 1 mm patch of red at the base of the hindwing in some individuals. This is in stark contrast to wild-type or BWPK butterflies which have black scales covering most of the body. Unlike the other BWPK butterflies, ivory homozygote butterflies do not fly and have not been observed to successfully mate or lay eggs. The wings appear to be of a normal strength and sturdiness, but flight is weak and will not occur when prompted by dropping (S1 Movie).

In order to investigate the genetic basis of these wing patterns, we sequenced 37 butterflies from the UT Austin *H. m*. BWPK stock to an average coverage of 13 x, including 11 Dark BWPKs, 9 Pale BWPKs and 10 ivory butterflies (Fig 1A).

### Autozygosity mapping identifies associated SNPs

We used a combination of GWAS and patterns of SNP autozygosity to determine the locations of regions associated with Mendelian pattern variants (Fig 2). As proof of principle, we mapped the Dennis pattern element which was previoulsy described as a *cis-*regulatory element of the gene *optix*. GWAS for presence vs absence of Dennis gave a narrow association peak centred near *optix*, in a region previously identified as associated with the Dennis pattern element (38) (S1 Fig). We then examined the inheritance patterns of SNPs, filtering for SNPs where butterflies with no Dennis element were homozygous for the reference allele, and individuals with the Dennis element were either heterozygous or homozygous for an alternate allele. In total, 3430 SNPs matched this pattern of inheritance, with 3425 in a cluster centred on the gene *optix*, indicating that autozygosity mapping is appropriate for mapping pattern elements in this data set.

**Fig 2:**
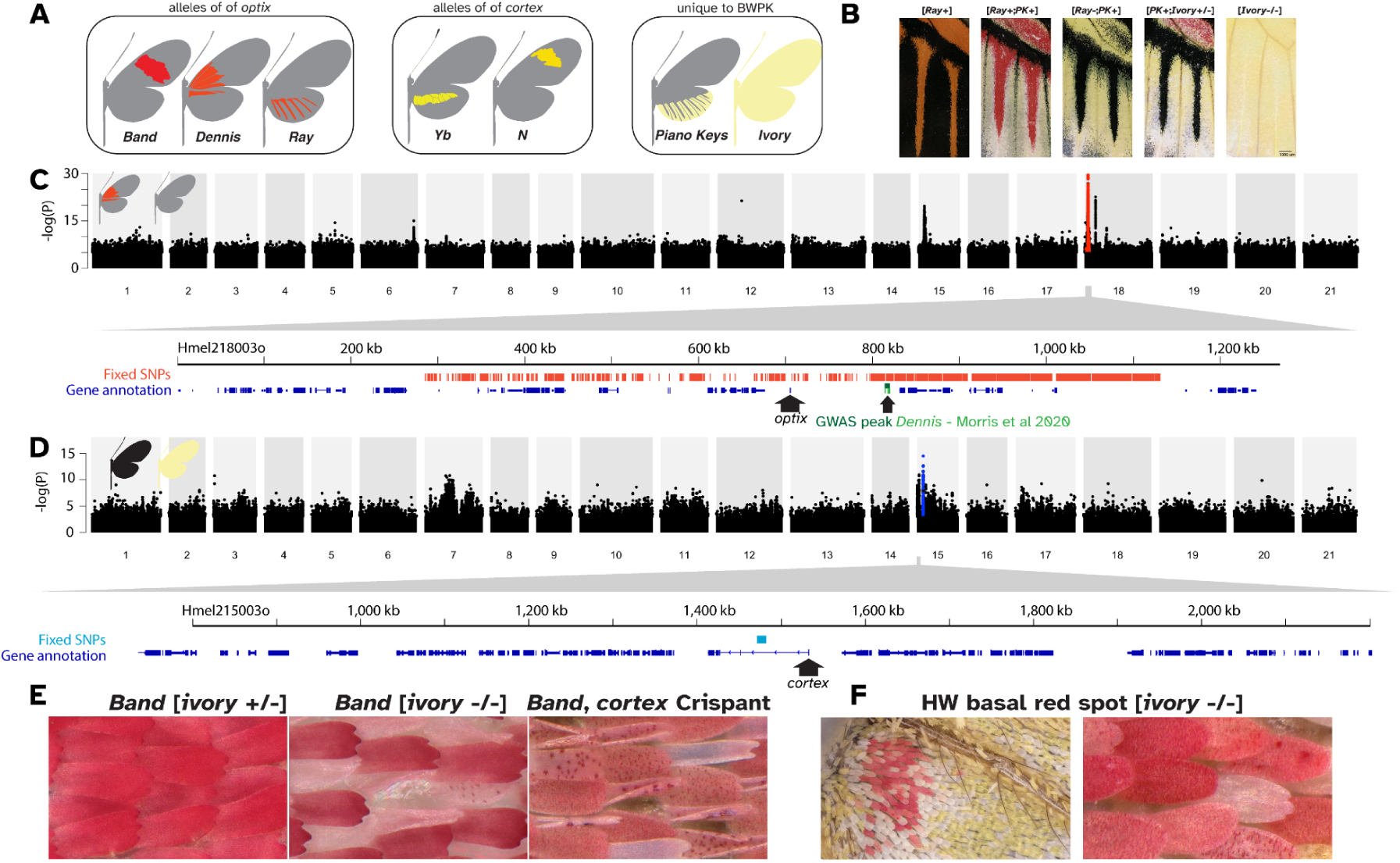
Genome wide association and autozygosity mapping in *H. melpomene* BWPK. **(A)** Cartoons of the allelic variation present in the *H. m*. BWPK stock. **(B)** the hindwing veins, showing the composite effects of different alleles depicted in panel A. **(C)** GWAS for the Dennis element; Manhattan plot of Wald test P-values, with fixed SNPs in red, and a magnified annotation of the region around the gene *optix*. **(D)** GWAS for ivory, with fixed SNPs in blue and a magnified annotation of the *cortex* region, with the point of the arrow at the annotated TSS. **(E)** comparison of red scales in [*ivory WT/WT*], [*ivory -/-*], and *cortex* crispant wings. **(F)** Some ivory butterflies have a small red dot at the base of the ventral hindwing, the only scales on the entire butterfly that are not white or yellow. They share the atypical phenotype of red scales from *cortex* crispants (**E**).

GWAS for ivory gave a broad association peak on chromosome 15 (Fig 2D). We expected dark morph BWPK butterflies to carry the reference allele, pale morphs to be heterozygous, and ivory morphs to be homozygous for a non-reference allele. We expected that at the causative locus, this allelic segregation pattern should be fixed in every individual. This was the case at just 131 SNPs in the whole genome, all within a 9 kb interval on chromosome 15, within the broad association peak (Fig 2D, blue marks). These SNPs sit within the first intron of the gene *cortex. Cortex* is a switch gene necessary for differentiation into black Type II and red Type III scales (41,42), and has previously been linked to melanic switches in many species, including in industrial melanism in the peppered moth (54,55).

We noted that in [*ivory WT/-*] butterflies, scales in red regions exhibited aberrant morphologies (Fig 2E-F). Granular clumps of red pigments could be observed in the body of some scales, whereas wild type scales are solidly-pigmented with no granularity. Additionally, we observed scales that were curled at the edges rather than flat. Similar scale phenotypes occur in the wings of *cortex* crispant butterflies (Fig 2E-F), thus supporting a role for *cortex* loss-of-function in the ivory colour phenotypes (42).

### Ivory butterflies carry a large deletion including the *cortex* promoter

Many phenotypes involve structural genomic variation, both under artificial selection (56), and in natural adaptation (57,58). As such, we checked read depth across this region in all our sequenced samples. Immediately adjacent to the block of fixed SNPs identified by autozygosity mapping, we found a large region where read depth in [*ivory -/-*] butterflies dropped to zero, while in [*ivory WT/-*] it dropped by half. This indicated that a large deletion spans the first exon and promoter of *cortex* (Fig 3A).

**Fig 3:**
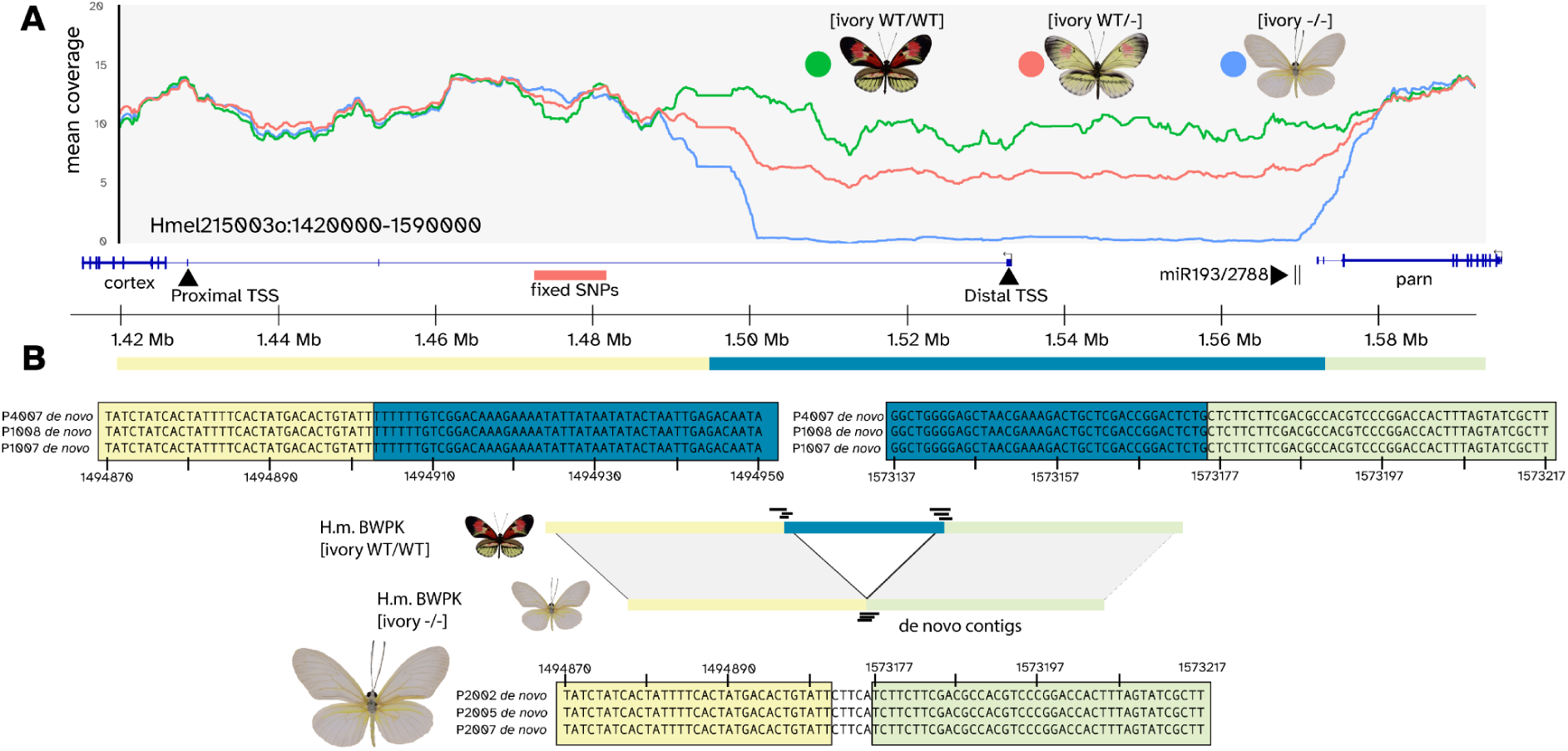
ivory is associated with a large deletion in the gene *cortex*. **(A)** mean coverage plot for the region around *cortex*, with [*ivory WT/WT*] in green, [*ivory WT/-*] in red, and [*ivory -/-*] in blue. **(B)** Alignments of *de novo* scaffolds, indicating the precise breakpoints of the *ivory* deletion. Colours indicate the orientation of sequences relative to the reference scaffold depicted in panel A.

To determine the precise breakpoints of this deletion, we *de novo* assembled short read data for all [*ivory WT/WT*] and [*ivory -/-*] individuals, and BLASTed the resulting contigs against the *cortex* region. This allowed us to recover individual contigs from some individuals that bridged the two ends of the deletion, giving a precise breakpoint at positions Hmel215003o:1494902-1573177, with a length of 78,275 bp relative to the reference genome (Fig 3B).

### The *ivory* deletion includes one of the two *cortex* transcription start sites

The *ivory* deletion contains the only annotated transcription start site (TSS) and promoter for *cortex*. While butterflies carrying this mutation can complete development, even with effects in scale development across the whole organism and the inability to fly, coding knockouts of *cortex* (including clonal G_0_ crispants) have been observed to have high levels of embryonic lethality, with very few crispant clones observed (42).

The ability of *ivory* mutants to survive in spite of the loss of the promoter, as well as the presence of a long first intron in *cortex*, led us to hypothesise that there may be a second, alternate TSS. In order to determine if this is the case, we mapped RNAseq from *H. melpomene* for a variety of samples including larval and pupal wings (59), adult ovaries (43), and embryos, adult head and adult abdomen (60), and looked at splicing and alignment of the 5’ end of the transcript. No expression was detected in adult head or abdomen, but in embryos, ovaries, and pupal wings, transcription initiates at position Hmel215003o:1424680 in the third annotated exon, which is adjacent to the coding sequence (the ‘proximal promoter’) (Fig 4A, S2-3 Figs). Expression in larval wings initiates at the annotated TSS (the ‘distal promoter’), and the exon with the proximal promoter is skipped. In pupal wings, transcription again initiated at the proximal promoter. This indicates that *cortex* has two alternative TSSs, a proximal promoter used in multiple tissues and a distal promoter specific to the larval wing. Only the distal promoter is included in the *ivory* deletion.

**Fig 4:**
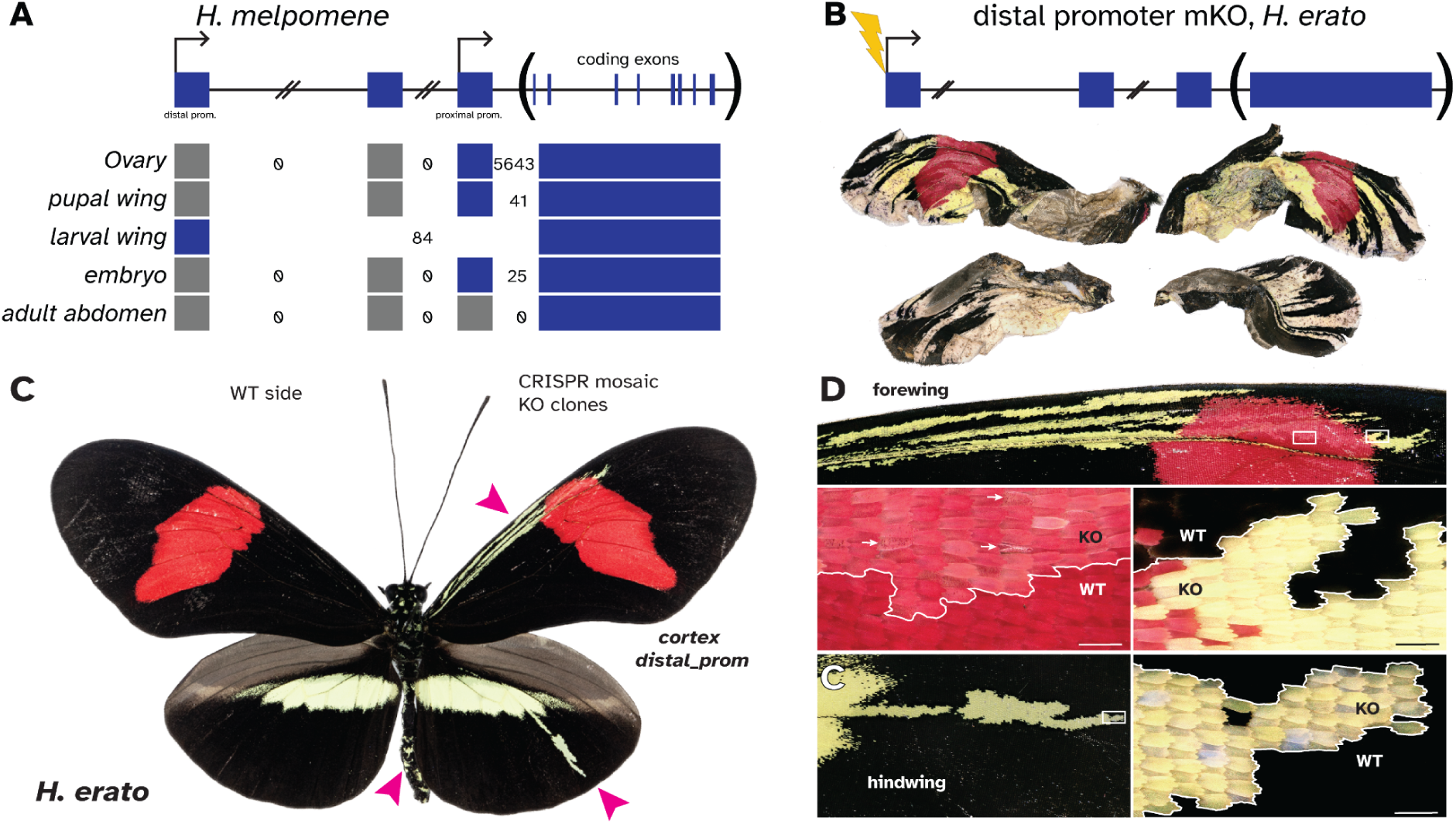
Knockout of the distal promoter phenocopies *cortex* protein coding knockouts. **(A)** mapping of RNAseq reads at *cortex* suggests that there are two promoters and transcription start sites. Numbers of intron-spanning reads are indicated (see Sashimi plots in Figs S2-3). **(B-C)** Knockout of the distal promoter in *H. erato* causes a transformation of black scales to white/yellow in a phenocopy of *cortex* protein coding knockouts. B shows a butterfly with extensive clones but that emerged poorly, indicating pleiotropic effects not observed in the *ivory* mutants, while C shows much smaller clones similar to those reported by Livraghi *et al*. 2021, with clone positions indicated by pink arrows. **(D)** High magnification images of mutant clones. Scale bars: 100 μm.

**Figure 5:**
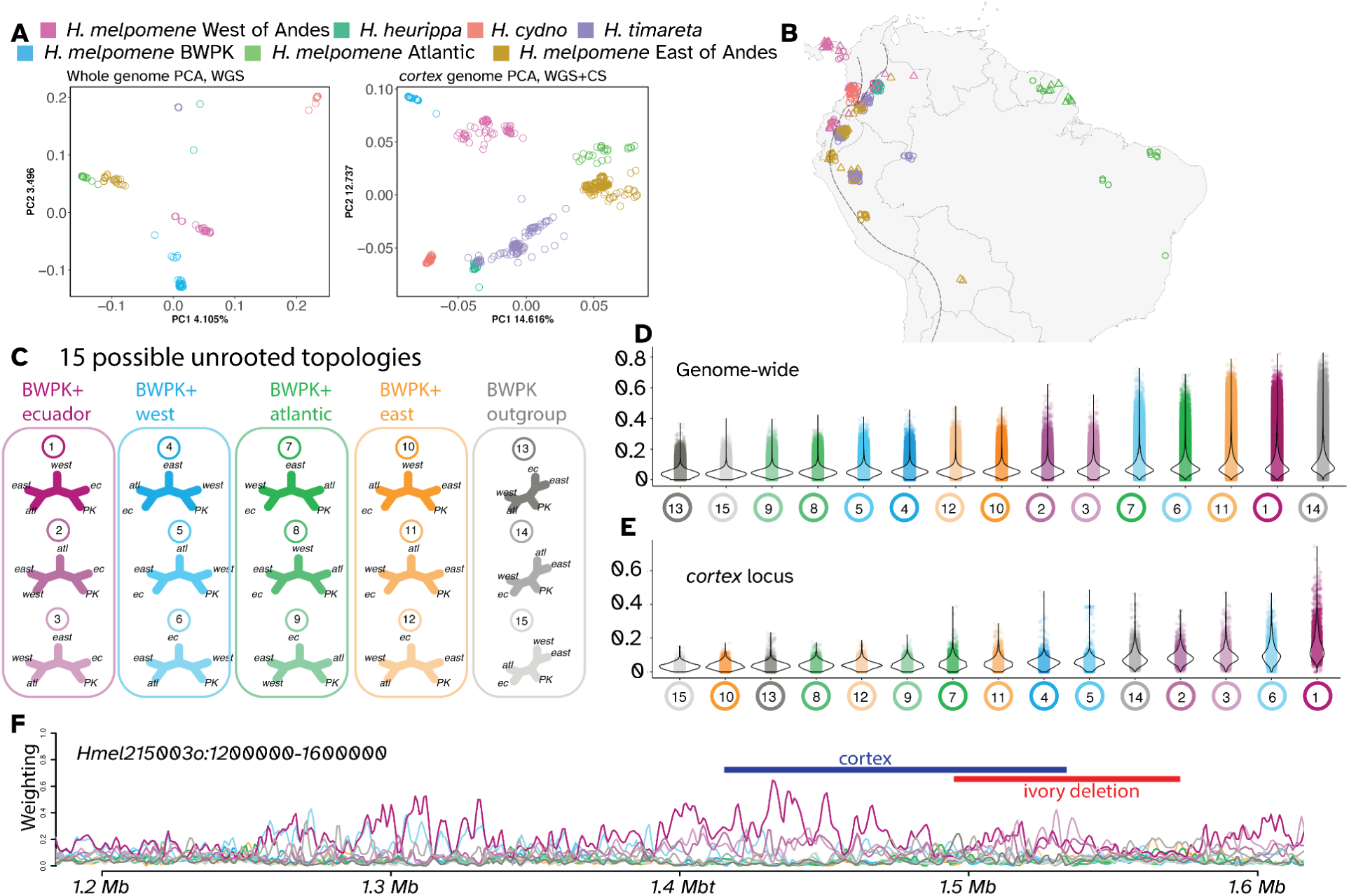
Phylogenetic clustering and weighting implies a mixed origin for *H. melpomene* BWPK. Clustering by PCA from both whole genome data (A) and from the *cortex* locus using additional selective sequencing data (B) show similar geographic clustering to that previously reported by Martin *et al*. (62), C), with the addition of separation between *H. melpomene* from East of the Andes and *H. melpomene* from French Guiana, Suriname and Brazil (here termed “Atlantic”). *H. m*. BWPK form a distinct cluster, closest to the West cluster of *H. melpomene*. Topology weighting was performed using TWISST (50), with five defined groups (West of Andes, Ecuador, East of Andes, Atlantic, and BWPK) (C). (D) Genome-wide, the most common topologies were 14 (BWPK as outgroup), 11 (BWPK clustered with East of Andes) and 1 (BWPK clustered with Ecuador). In contrast to this in (E), TWISST results at the *cortex* locus showed that topology 1 was most common, supporting an Ecuadorian origin for the *cortex* variants in *H. m. BWPK*.

### Mutagenesis of the distal promoter phenocopies *cortex* null effects

We reasoned that if the loss of the distal promoter causes the ivory phenotype, knocking out the promoter with CRISPR/Cas9 should phenocopy the effects of the *ivory* deletion. We generated G_0_ mosaic knockouts (crispants) of the promoter in the related species *Heliconius erato* (Fig 4B-D). Resulting butterflies had extensive yellow/white clones on the wings, however several failed to emerge from the pupa, and a low survival rate and penetrance were observed (Table 1), suggesting that the few wing crispants obtained are rare mosaic escapers of a loss-of-function experiment with lethal effects (61). Thus while the pleiotropic effects of deleting the TSS were reminiscent of the previously reported *cortex* protein coding knockouts, which also exhibited low survival and penetrance (42), this experiment did not fully recapitulate the ability of the *ivory* deletion allele to generate large wing clones. This difference in pleiotropy may be due to the use of *H. erato* in the CRISPR assay, and further experiments are needed to test the functionality of deleted elements within a *H. melpomene* stock. However, we conclude from these experiments that the disruption of 5’ distal elements of *cortex* are necessary for normal color scale patterning, consistent with a causal role of the 78 kb deletion in underlying the ivory phenotypes.

### Demographic origins

The *H. m*. BWPK stock was kept in insectaries for 30 years (c. 400 generations) before producing the *ivory* mutation. The precise history and ancestry of the stock is not recorded, but we do know that the original stock was generated with a mix of butterflies from multiple locations. after the addition of butterflies from the vicinity of Tinalandia, Ecuador in the 1980s. The first pale PK (*i*.*e*. heterozygotes for *ivory*) appeared in the PK culture at Butterfly World after 2013, and were shared with L.E.G. for study in 2014-15.

Whole genome PCA clustering of a geographic spread of samples of *H. melpomene*, as well as sister taxa *H. cydno* and *H. timareta*, replicates the geographic and species clustering previously described by Martin *et al*. (62), with the addition of a distinct “Atlantic” group within *H. melpomene* (S1 Table). *H*.*m*. BWPK cluster together near *H. melpomene* from the west of the Andes. Similarly, PCA of just the *cortex* locus, using many more individuals generated by selective sequencing by Moest et al (63) (S3 Table), also places BWPK closest to western *H. melpomene*.

Given the mixed ancestry of *H. m* BWPK, and the observation that ivory patterns emerged after introduction of butterflies from Ecuador, we examined heterogeneity in genome-wide ancestry by building phylogenetic trees in windows of 50 SNPs across the genome and using topology weighting to determine the closest neighbour of *H*.*m* BWPK at all genomic positions. Of the 15 possible topologies, the most frequent topology places BWPK as an outgroup to the four other *H. melpomene* taxa. The second most common topology groups BWPK with *H. melpomene* from western Ecuador (*H. m vulcanus* and *H. m. plesseni*), and the third most common places BWPK with *H. melpomene* from the East of the Andes. In contrast, sliding windows across the *cortex* locus (including the selective sequencing data) show the most common tree topology here places BWPK sister to the western Ecuador taxon. This result is supportive of *H*.*m*. BWPK having an admixed genome, but with the allele at *cortex* having originated in Ecuador. Importantly, the *ivory* deletion was not detected in any of 458 wild individuals.

## Discussion

Using whole genome sequencing of a set of related individuals, we were able to map the *ivory* mutation to a large deletion that removed the distal promoter of the wing patterning gene *cortex*. Cortex was first identified as a switch between melanic and non-melanic patterns in both *Heliconius* and the peppered moth *Biston betularia*, and has later been mapped as a switch between melanic and non-melanic scale types in a number of other species, including three other geometrid moths (55), and *Papilio* butterflies (64), and also has a role in wing pattern polyphenism in *J. coenia* (65). The list of genes that have been identified as the targets of selection in butterfly wing pattern variation includes transcription factors like Optix and Bab (66,67), signalling pathway components like WntA (68), or terminal effector genes like Yellow (69), all of which have molecular functions that support a role in developmental patterning or pigment synthesis (34). Cortex, on the other hand, has no clear functional association with colour pattern or cell differentiation, and yet has been mapped in a very broad phylogenetic spread of species, suggesting its function may be ancestral in the Lepidoptera. The gene does not appear to have a role in *Drosophila* wing development or patterning (41), limiting our ability to make inferences about its molecular function and interactions. In identifying this large deletion, this study provides novel insight into the genetics of this hotspot locus, and highlights the potential utility of looking at insectary mutants to understand butterfly wing development.

The loss of all black and red scales in the *ivory* mutant supports a model in which white/yellow scales are the ‘default’ state during scale cell differentiation, with scale cell precursors failing to differentiate into red or black cell types. As such, when comparing pattern homologies between butterflies, *Heliconius* white/yellow regions can be considered as the ‘background’, and black and red regions can be considered as ‘pattern elements’ layered on top of this background. This model was initially proposed based on observation and ultrastructure and pigmentation of scales (70), as well as on the analysis of pattern homologies of the Heliconiini (71).

### Structural variation and mutations

Large deletions have repeatedly been observed in cases of both natural variation and domestication, with 29 deletions affecting regulatory regions which are larger than 1 kb listed on GepheBase (72). Recurrent deletions of a *pitx1* enhancer in sticklebacks, up to 8 kb in length, have caused convergent pelvic reduction (73), and the deletion of a 60.7 kb regulatory region at the *AR* gene in humans, which removes enhancers present in chimpanzee and conserved in other mammals, leads to loss of penile spines (74). Similarly large deletions have also been confirmed in the genetic basis of domestication phenotypes, including a 44 kb deletion, also at *pitx1*, causing feathered feet in chickens (75), and a 141 kb deletion upstream of *agouti/ASIP* in Japanese quail (76).

The large deletion that causes ivory is a form of structural variation. Other structural variants affecting this locus have been found in wild populations of *Heliconius*; multiple inversions have formed a wing pattern supergene in *H. numata*, and an independent inversion, similar in size and position, has been found in multiple species of the *erato* clade (77,78). It is possible that this genomic region will prove to be prone to a higher-than-average level of structural variation, or, conversely, that the region is more tolerant to structural variation than other parts of the genome.

### Regulatory consequences of the *ivory* mutation

Large deletions have the potential to remove a large section of *cis-*regulatory sequence, as well as transcribed sequences. The *ivory* deletion causes the loss of one of two promoters. As many as 40% of developmentally expressed genes in *Drosophila* have two promoters which can cause distinct regulatory programs (79), and multiple promoters are also common in human genes (80). This both increases the complexity of gene regulatory interactions and increases the number of transcript isoforms per gene - for example, an alternate promoter in the *Drosophila* gene *Zfh1* creates an isoform that has a shorter 5’ UTR which is missing miRNA seed sites on its 5’UTR, permitting differential degradation of the mRNA by *miR-8* (81). We determined that the *ivory* deletion contains one annotated promoter of *cortex*, but that in some contexts during development, another transcription start site is used, much closer to the translation start site. Alternate promoter usage is likely to be common and widespread in animals.

We expect that the use of two separate, context-specific promoters at *cortex* will create a hierarchical regulatory organisation, where CREs used in different tissues or at different times can loop to different promoters. Additionally, transcripts from each of the two promoters contain a different arrangement of 5’ non-coding exons, giving them different UTRs, possibly leading to differential translational regulation or degradation of the mRNA.

The *ivory* deletion does not include protein-coding sequence, which led us to hypothesise that by removing just one promoter, *cortex* would still be expressed in tissues that use the alternate promoter, and that therefore *ivory* mutants would bypass the pleiotropic, highly lethal effects observed in CRISPR experiments that target coding exons of *cortex* (42). However, crispants with deletions in the distal promoter in *H. erato* did not appear to be free of these pleiotropic effects, and exhibited high lethality and comparatively small clone sizes, dissimilar to the *H. melpomene* [*ivory -/-*] mutants. This may be due to inherent differences in the regulatory function of the distal TSS between these two species, or to the existence of other functional sequences within the deletion which might be necessary for generating the pale phenotypes. Specifically, the 78-kb deletion removes part of the 3’UTR of the neighbouring gene *parn* as well as two miRNAs (82), and it is also possible that a large deletion affects other cortex cis-regulatory elemnts or affect 3D chromatin interactions on a large portion of the chromosome. Additionally, the *ivory* butterflies carry a set of fixed SNPs adjacent to the deletion, which could contribute to the phenotype. In summary, the complete loss of black and red scales caused by the *ivory* deletion and by CRISPR-Cas9 mutagenesis in two *Heliconius* species indicates that either the distal promoter or another functional sequence within the 5’ non-coding region of *cortex* are necessary for black and red scale development.

### Heterozygote advantage of deleterious mutations in captivity

The *cortex* mutant phenotype has been artificially selected by a breeder in the heterozygous state, when unusually widespread yellow and white patterns emerged in the stock. The heterozygote carriers of the *ivory* mutation ([*ivory WT/-*]) do not appear to have any reduction in fitness or vigor. In contrast, homozygous *ivory* butterflies of either sex are unable to fly, limiting their ability to feed and reproduce – we have not observed any successful matings involving an [*ivory -/-*] butterfly. Those recessive effects make the mutant allele unfit for breeding in the homozygous state, resulting in a form of heterozygote advantage where mutant heterozygotes are selectively bred in spite of inviable or undesirable effects in homozygotes(83,84). We compiled similar situations from the literature using a database of gene-to-phenotype relationships (72), and found 38 analogous gene-to-trait relationships spanning domesticated mammals and birds (Table 2). Large structural variations (SVs) such as the *ivory* deletion accounted for 9 out of 46 derived alleles in this dataset, suggesting that the recessive deleterious effects of macromutations occasionally provide heterozygous states of interest to artificial selection. Of note, 17 out the 38 cases of captive heterozygote advantage involve selection for depigmentation traits, highlighting the trend among breeders and fanciers to select for conspicuous variants (Table 2). Finally, cases of heterozygous advantage also occur in the wild, such as in ruff birds where a large inversion haplotype provides male coloration and behavioral traits that are maintained by sexual selection in spite of being recessive lethal (85). In wild populations of the polymorphic butterfly *Heliconius numata, cortex* itself is situated in the center of an inversion polymorphism under balancing selection (86). Further work will be needed to determine the sub-gene level elements that are within the *ivory* deletion and mediate homozygous inviability.

**Table 2:** Known cases of heterozygote advantage in animal breeding.

| Gene | Species | Heterozygote advantage (HA) trait for breeding | Detrimental recessive effect | Nb of HA alleles | Mutation types |
| --- | --- | --- | --- | --- | --- |
| *ACAN* | Cattle | Short stature | Lethal | 1<br>1 | Frameshift (4-bp ins)<br>New start codon (SNP) |
| *ACAN* | Horse | Short stature | Lethal | 1<br>1 | Loss of 6 a.a. (18-bp del)<br>Frameshift (1-bp del) |
| *ALX1* | Cat | Brachycephaly | Frontonasal dysplasia | 1 | Loss of 4 a.a. (12-bp del) |
| *BBS9* | Pig | Fast growth | Lethal (misexpressed *BMPER*) | 1 | Large SV (212-kb del) |
| *BMP15* | Sheep | Increased female fecundity | Female infertility | 5<br>2<br>1<br>1<br>1 | Missense (SNP)<br>Nonsense (SNP)<br>Frameshift (1-bp ins)<br>Frameshift (17-bp del)<br>Cis-regulatory (SNP) |
| *cortex* | *Heliconius* | White patterns | Flightless | 1 | Large SV (78-kb del) |
| *EDNRB* | Horse | White patterns | Lethal | 1 | Missense (SNP) |
| *FGF3/4/19* cluster | Dog | Dorsal hair ridge | Dermoid sinuses | 1 | Large SV (133-kb duplication) |
| *FGFR3* | Sheep | Increased bone length | Chondrodysplasia | 1 | Missense (SNP) |
| *FOXI3* | Dog | Hairlessness | Ectodermal dysplasia | 1 | Frameshift (7-bp duplication) |
| *GDF9* | Sheep | Increased female fecundity | Female infertility | 5 | Missense (SNP) |
| *HMGA2* | Rabbit | Short stature | Lethal | 1 | Large SV (12.1-kb del) |
| *HOXC10* | Chicken | Head crest | Cerebral Hernia - likely epistatic | 1 | Large SV (197-bp duplication) |
| *IHH* | Chicken | Short legs | Lethal | 1 | Large SV (11.9-kb del) |
| *KIT* | Camel | White patterns | Lethal | 1 | Frameshift (1-bp del) |
| *KIT* | Dog | White patterns | Lethal | 1 | Frameshift (1-bp ins) |
| *KIT* | Donkey | White patterns | Lethal - not proven | 1<br>1 | Missense (SNP)<br>Splice Site mutation (SNP) |
| *KIT* | Fox | White patterns | Lethal | 1<br>1 | Missense (SNP)<br>Splice Site mutation (SNP) |
| *KIT* | Horse | White patterns | Lethal | 3<br>2<br>2<br>1<br>1 | Frameshift (1-bp, 1-bp, 4-bp del)<br>Large SV (1.3-kb, 1.9-kb del)<br>Nonsense (SNP)<br>In-Frame Del (54-bp)<br>Splice Site mutation (SNP) |
| *KIT* | Pig | White patterns | Lethal - not proven | 1 | Splice Site mutation (SNP) |
| *MITF* | Buffalo | White patterns | Microphthalmia (inferred from mice) | 1<br>1 | Nonsense (SNP)<br>Splice Site mutation (SNP) |
| *MITF* | Dog | White patterns | Deafness (risk increase*) | 1 | Cis-regulatory (195-bp ins) |
| *MITF* | Horse | White patterns | Microphthalmia (inferred from mice) | 2<br>1<br>1 | Large SV (8.7-kb, 63-kb del)<br>Frameshift (4-bp del)<br>Missense (SNP) |
| *MITF* | Quail | White patterns | Small size and slow growth | 1 | Frameshift (2-bp del) |
| *Mlana* | Pigeon | White patterns | Eye defects | 4 | CNV including 3 other genes |
| *MNR2* locus | Chicken | Head comb | Male infertility | 1 | Large SV (7.4-Mb inversion) |
| *MRC2* | Cattle | High muscle | Musculoskeletal defects | 1<br>1 | Frameshift (2-bp del)<br>Missense (SNP) |
| *MSTN* ** | Dog | Racing performance | Non-conform appearance | 1 | Frameshift (2-bp del) |
| *PAX3* | Horse | White patterns | Lethal | 2 | Missense (SNP) |
| *PMEL17* | Dog | White patterns | Auditory and ocular defects | 6 | Splice Site mutation (TE insertions) |
| *PMEL17* | Horse | White patterns | Ocular defects | 1 | Missense (SNP) |
| *RNASEH2Ba* locus | Cattle | Milk yield (causal gene unknown) | Lethal (due to *RNASEH2Ba* KO) | 1 | Large SV (660-kb deletion) |
| *RYR1* | Pig | Meat yield and quality | Hyperthermia | 1 | Missense (SNP) |
| *PRLR* / *SPEF2* | Pig | Increased female fecundity | Sperm defects (due to *SPEF2* KO) | 1 | Cis-regulatory (TE insertion) |
| *SPINT1* | Gecko | White patterns | Metastases | 1 | Unknown |
| *STX17* | Horse | White patterns | Melanoma | 1 | Cis-regulatory (4.6-kb duplication) |
| *TBXT* (Brachyury) | Cat | Short or no tail | Lethal | 4 | Frameshift (1-bp del; 14-bp ins) |
| *TBXT* (Brachyury) | Dog | Short or no tail | Lethal | 1 | Missense (SNP) |
| *TRPM1* | Horse | White patterns | Ocular defects | 1 | Cis-regulatory (TE insertion) |
| *TRPV4* | Cat | Short ears | Chondrodysplasia | 1 | Missense (SNP) |

\* See Gephebase (www.gephebase.org) for references. The corresponding entries can be retrieved using a search for "@HeterozygoteAdvantage".
\*\* recommendations to outcross are recent in the affected stock.
\*\*\* *MSTN* loss-of-function alleles exist in other bred species. Homozygotes sometimes requires assisted birthing, but are viable and artificially selected.

## Conclusion

By using autozygosity mapping and association on a comparatively small pool of individuals, we were able to identify a structural mutation involved with butterfly wing pattern. This approach may prove fruitful in other studies of butterfly wing patterning or in the identification of *de novo* mutants, especially if combined with recent developments in whole genome sequencing from dried museum specimens (87,88). This could allow the mapping of other cases like *Hindsight* in *Junonia coenia*, or *pseudozorro* in *Parnassius apollo* (89,90). The identification and characterisation of spontaneous mutants has provided very valuable insights into the genetics of development in model organisms. Further whole genome sequencing of deleterious mutations in butterflies will help to identify parts of the genome that are functionally required for normal development, which will assist in a more complete understanding of their evolution and development.

## Data availability

Whole genome sequences are accessible on the NCBI SRA under the Bioproject accession PRJNA610063.

## Acknowledgements

We thank Ron Boender of Butterfly World, FL for generously providing living specimens for study by the Gilbert Lab at UT Austin. Thanks to greenhouse technicians T. Freiburger, N. Fogel, I. Terry, and E. Reese for keeping plant and butterfly cultures healthy in the Austin facility during the study, and G. Julian at the University of Cambridge for expert stock maintenance of *H. erato*. We also thank the GWU HPC team for computing infrastructure (91), and S.M. Van Belleghem for comments on an earlier draft.

---

**S1 Table :** Samples and accessions.

| Accession | BioSample<br>Accession | Sample<br>Name | Phenotype | Inferred Genotype | Date<br>Collected | Coverage | %GC |
| --- | --- | --- | --- | --- | --- | --- | --- |
| SRR11243376 | SAMN14276602 | P1001 | Piano Keys, dark | *B/-, d/d, r/r, [ivory WT/WT]* | 30 Oct 2014 | 13.01 | 32.53 |
| SRR11243375 | SAMN14276603 | P1002 | Piano Keys, dark | *B/-, d/d, r/r, [ivory WT/WT]* | 30 Oct 2014 | 14.94 | 32.97 |
| SRR11243364 | SAMN14276604 | P1006 | Piano Keys, dark | *B/-, d/d, r/r, [ivory WT/WT]* | 23 Feb 2014 | 16.18 | 32.52 |
| SRR11243353 | SAMN14276605 | P1007 | Piano Keys, dark | *B/-, D/-, r/r, [ivory WT/WT]* | 19 Feb 2013 | 14.93 | 32.38 |
| SRR11243344 | SAMN14276606 | P1008 | Piano Keys, dark | *B/-, D/d, r/r, [ivory WT/WT]* | 19 Feb 2013 | 13.42 | 32.83 |
| SRR11243343 | SAMN14276607 | P1009 | *cydno galanthus* |  | 19 Nov 2019 | 16.14 | 32.41 |
| SRR11243342 | SAMN14276608 | P1010 | *cydno* x pale PK |  | 19 Nov 2019 | 14.07 | 32.37 |
| SRR11243341 | SAMN14276609 | P1011 | F1 pale PK x *cydno* |  | 6 Feb 2019 | 16.14 | 32.5 |
| SRR11243340 | SAMN14276610 | P1012 | F1 pale PK x *cydno* |  | 2 Feb 2019 | 16.62 | 32.6 |
| SRR11243339 | SAMN14276611 | P1013 | BC of F1 x *cydno* |  | 4 Feb 2019 | 12.49 | 32.66 |
| SRR11243374 | SAMN14276612 | P1014 | BC of F1 x *cydno* |  | 4 Feb 2019 | 14.77 | 32.19 |
| SRR11243373 | SAMN14276613 | P1015 | BC of F1 x *cydno* |  | 19 Feb 2019 | 15.93 | 32.44 |
| SRR11243372 | SAMN14276614 | P2001 | ivory | *[ivory -/-]* | 21 Jan 2019 | 15.37 | 32.24 |
| SRR11243371 | SAMN14276615 | P2002 | ivory | *[ivory -/-]* | 21 Jan 2019 | 11.69 | 32.48 |
| SRR11243370 | SAMN14276616 | P2003 | ivory | *[ivory -/-]* | 6 Jan 2019 | 8.95 | 32.45 |
| SRR11243369 | SAMN14276617 | P2004 | ivory | *[ivory -/-]* | 30 Sep 2019 | 16.11 | 32.49 |
| SRR11243368 | SAMN14276618 | P2005 | ivory | *[ivory -/-]* | 23 Sep 2019 | 16.19 | 32.74 |
| SRR11243367 | SAMN14276619 | P2006 | ivory | *[ivory -/-]* | 10 Feb 2019 | 11.47 | 33.03 |
| SRR11243366 | SAMN14276620 | P2007 | ivory | *[ivory -/-]* | 21 Jan 2019 | 15.44 | 32.77 |
| SRR11243365 | SAMN14276621 | P2008 | ivory | *[ivory -/-]* | 1 Oct 2019 | 17.40 | 32.6 |
| SRR11243363 | SAMN14276622 | P2009 | ivory | *[ivory -/-]* | 24 Jan 2019 | 12.99 | 32.64 |
| SRR11243362 | SAMN14276623 | P2010 | Piano Keys, pale | *B/-, d/d, r/r, [ivory WT/-]* | 24 Jan 2019 | 13.25 | 32.78 |
| SRR11243361 | SAMN14276624 | P2011 | ivory | *[ivory -/-]* | 20 Jun 2019 | 16.63 | 32.32 |
| SRR11243360 | SAMN14276625 | P3005 | Piano Keys, dark | *B/-, d/d, r/r, [ivory WT/WT]* | 19 Nov 2019 | 15.92 | 32.71 |
| SRR11243359 | SAMN14276626 | P3006 | Piano Keys, dark | *B/-, D/-, R/- [ivory WT/WT]* | 19 Nov 2019 | 14.44 | 32.71 |
| SRR11243358 | SAMN14276627 | P3007 | Piano Keys, dark | *B/-, D/-, R/- [ivory WT/WT]* | 16 Dec 2018 | 16.45 | 32.43 |
| SRR11243357 | SAMN14276628 | P3016 | Piano Keys, dark | *B/-, d/d, r/r, [ivory WT/WT]* | 27 Nov 2018 | 9.35 | 32.58 |
| SRR11243356 | SAMN14276629 | P3017 | Piano Keys, dark | *B/-, d/d, r/r, [ivory WT/WT]* | 14 Dec 2018 | 19.50 | 32.4 |
| SRR11243355 | SAMN14276630 | P3019 | Piano Keys, dark | *B/-, d/d, r/r, [ivory WT/WT]* | 27 Jan 2019 | 15.40 | 32.41 |
| SRR11243354 | SAMN14276631 | P4001 | Piano Keys, dark | *B/-, d/d, r/r, [ivory WT/-]* | 19 Feb 2019 | 13.24 | 32.89 |
| SRR11243352 | SAMN14276632 | P4002 | Piano Keys, pale | *B/-, D/-, R/-, [ivory WT/-]* | 19 Nov 2019 | 13.85 | 32.62 |
| SRR11243351 | SAMN14276633 | P4003 | Piano Keys, pale | *B/-, d/d, r/r, [ivory WT/-]* | 5 Feb 2019 | 13.20 | 32.48 |
| SRR11243350 | SAMN14276634 | P4004 | Piano Keys, pale | *B/-, d/d, r/r, [ivory WT/-]* | 7 Dec 2019 | 15.82 | 32.87 |
| SRR11243349 | SAMN14276635 | P4005 | Piano Keys, pale | *B/-, d/d, r/r, [ivory WT/-]* | 19 Nov 2019 | 17.01 | 32.68 |
| SRR11243348 | SAMN14276636 | P4006 | Piano Keys, pale | *B/-, D/-, R/-, [ivory WT/-]* | 19 Nov 2019 | 18.17 | 32.63 |
| SRR11243347 | SAMN14276637 | P4007 | Piano Keys, pale | *B/-, d/d, r/r, [ivory WT/-]* | 26 Jan 2019 | 9.74 | 32.77 |
| SRR11243346 | SAMN14276638 | P4008 | Piano Keys, pale | *B/-, d/d, r/r, [ivory WT/-]* | 19 Dec 2018 | 11.18 | 33.58 |
| SRR11243345 | SAMN14276639 | P4009 | Piano Keys, pale | *B/-, d/d, r/r, [ivory WT/-]* | 19 Dec 2019 | 13.96 | 32.63 |

**S1 Figure:**
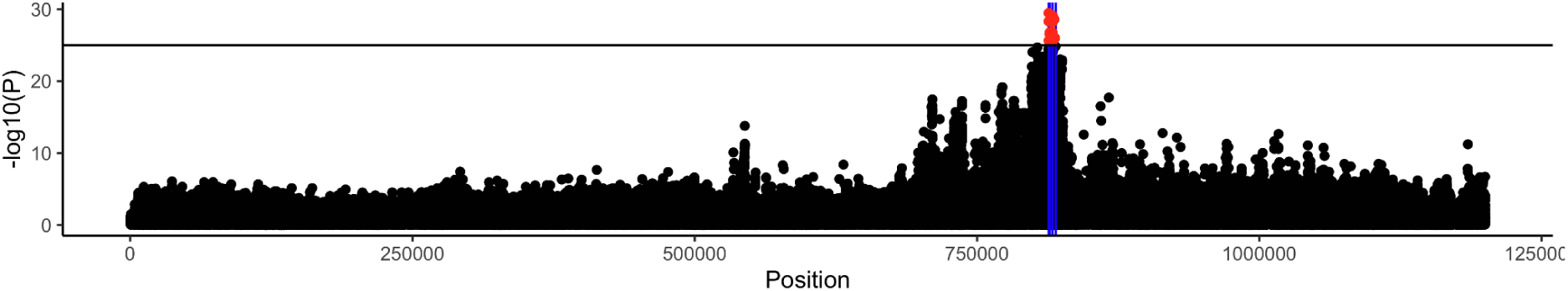
Dennis GWAS zoomed on Hmel218003o, showing the tip of the association peak (-log10P > 25) aligning precisely with the Dennis element designated by Morris et al (2020).

**S2 Figure:**
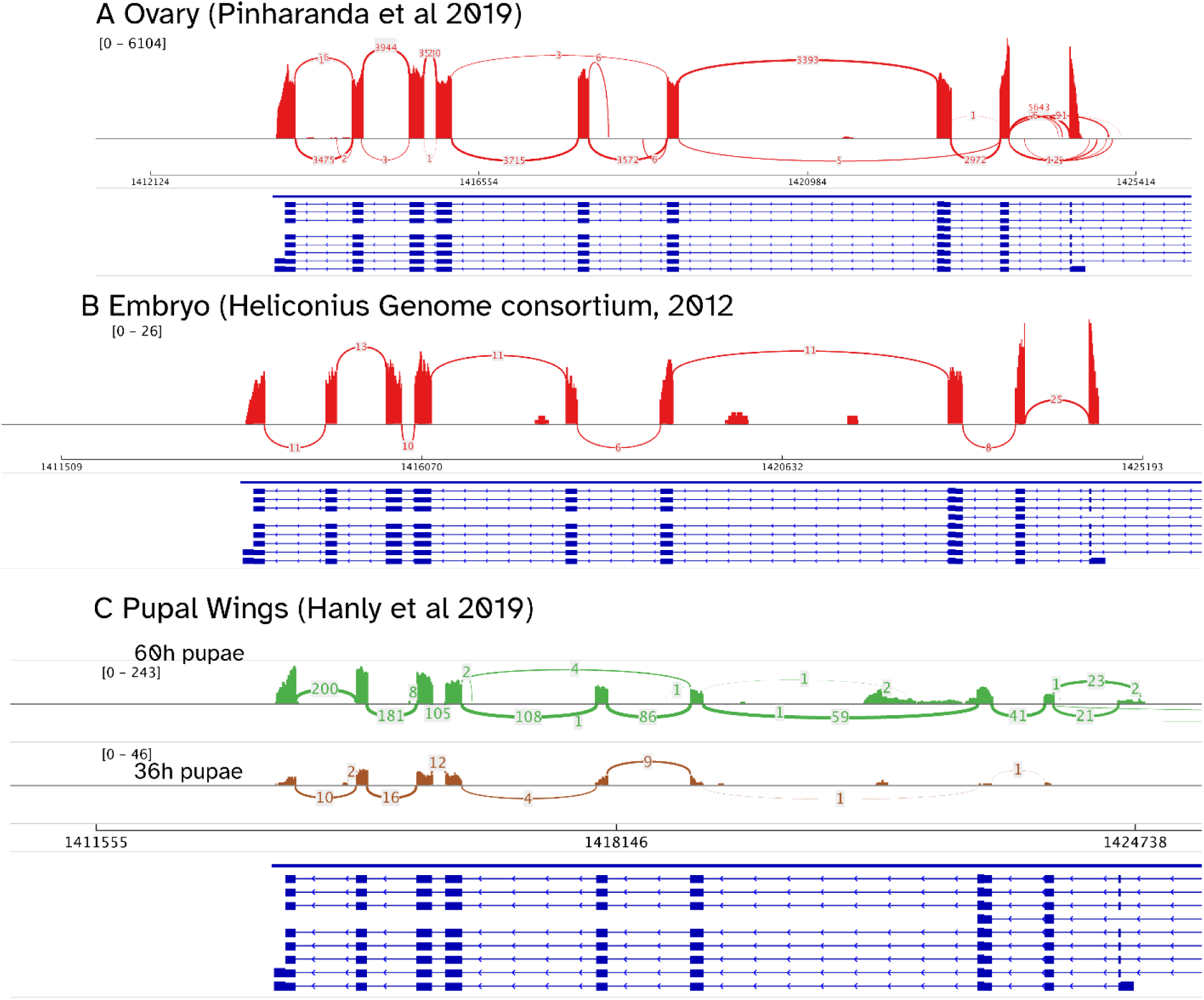
Sashimi plot of *H. melpomene* RNA splicing at *cortex*. Ovary from Pinharanda et al 2019 (A), embryo from Dasmahapatra et al 2012 (B), pupal wings from Hanly et al 2019 (C), with 60h pupal wings above and 36h pupal wings below. Annotations of splice variants from *cortex* from the Hmel2.5 annotation are included, running in the antiparallel direction. Note that while expression of cortex is at least 20-fold higher in ovary compared to embryo or wing, no expression or splicing to the distal TSS was detected.

**S3Figure:**
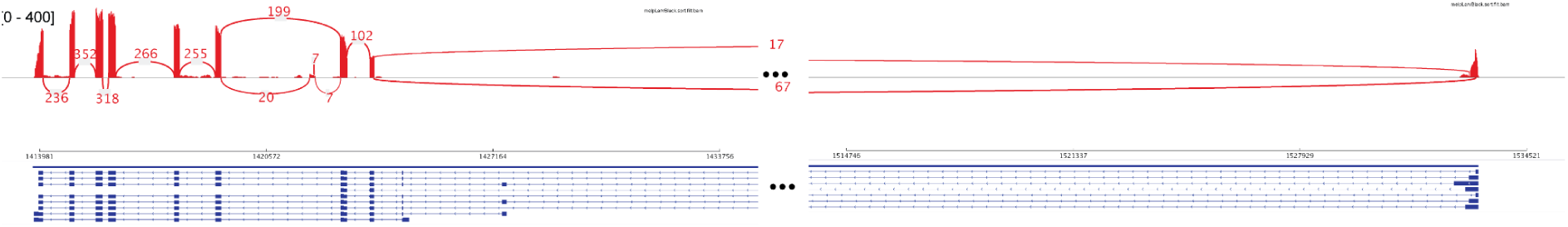
Sashimi plot of *H. melpomene* RNA splicing at the full cortex annotation from larval wings, from Hanly et al 2019. The ellipsis indicates an 81kb interval. Here, the TSS at the distal promoter is used, and the proximal promoter and TSS are not transcribed.

